# Reciprocal effective conductance-based inference of protein-protein relationships delineates protein network hubs and macromolecular properties

**DOI:** 10.64898/2026.10.12.758675

**Authors:** K C Rajendra, Sihang Zhou, Ruiying Cheng, Marco Andres Penaherrera, Simon Lizarazo, Kevin Van Bortle

## Abstract

Protein-protein interactions (PPIs) are fundamental to the regulation and function of all cellular processes, yet our current understanding of the human interactome is limited by the specific PPI-related experiments performed to date. Moreover, whereas proteins with minimal research interest may have relatively few interaction-probing studies, other factors such as cancer related proteins are frequently represented in PPI studies, producing a highly uneven bias across current PPI data. Here, we apply effective conductance – a concept from circuit theory – to infer proximity relationships across all individual protein pairs on the basis of 1.2 million reported human PPI observations. Using a reciprocal analysis framework built on effective conductance, we identify statistically meaningful interactions within the network while otherwise filtering nonrelevant associations that are abundant in proteins with strong popularity bias. While this approach helps clarify the structural and functional properties of both minimally and maximally connected proteins, we further compute an interaction index between all macromolecular, multi-subunit complexes based on underlying protein-protein conductance scores. Leveraging this framework, which we name CoBRAA (Conductance-Based Reciprocal Association Analysis), we highlight unexpected findings, such as evidence that DNA-dependent RNA polymerase I, II, and III are characterized by surprisingly pervasive cross-complex interactions between enzyme-specific subunits, suggesting a potential for crosstalk between all three RNA polymerases. We further introduce CoBRAA as an open-access research tool that enables custom, user-specified exploration of both protein- and macromolecular-level interactions. We expect this resource to facilitate broader investigation and hypothesis generation across a wide range of proteins, pathways, and macromolecular systems.

## Introduction

Protein-protein interactions (PPIs) coordinate nearly every cellular process, from signal transduction and subcellular compartmentalization to transcription regulation and the biosynthesis of proteins and macromolecular assemblies^1^. However, our understanding of the protein interactome in humans and other species remains both incomplete and uneven. Though experimental maps derived from yeast two-hybrid, affinity capture, complementation assays and others have helped to uncover thousands of context-linked PPIs, these data are far from comprehensive^2^. A second complication arises due to experimental bias. Proteins with long-standing biomedical interest, such as oncogenic and tumor suppressing transcription factors, are extensively profiled, whereas others remain comparatively unexplored^3^. This skew produces a distorted representation of the protein interactome, with some factors appearing highly connected to other proteins potentially due to study frequency^4,5^.

Innovative strategies that build on currently available experimental data are needed to address representation bias and better delineate the nature of the human protein interactome. To this end, we introduce a framework that considers how two proteins are positioned relative to one another within the broader network of ~1.2 million currently mapped PPIs. Here, we specifically apply the principle of effective conductance, a network-based measure of association that considers both direct and indirect paths (i.e., between two disconnected nodes)^6-10^, to gain a quantitative measure of distance between all protein-pairs within the human proteome. Using effective conductance in this way, we determine the reciprocal significance of connectivity between all protein pairs, facilitating the pruning of potentially spurious interactions driven by experimental bias, and the inference of meaningful protein-protein relationships within the network. Extending this framework to consider macromolecular relationships, we further quantify connectivity between large, multi-subunit complexes. We demonstrate the utility of this framework in the form of ‘CoBRAA’ - an interactive research tool that gives users the opportunity to explore both protein- and macromolecular-level analyses derived from conductance-based reciprocal association analysis.

## Results

### Effective conductance-based modeling of all human protein-protein interactions and relationships

To establish a human protein-protein interaction network, we first retrieved all currently available PPIs from the BioGRID database^11,12^, restricting for experimentally observed physical interactions (see methods). In total, 1.2 million PPIs were transformed into a PPI network, such that each edge represents the number of independent PPI observations between each pair of protein nodes (Figure 1a). By conceptualizing PPI frequency as the conductance within an electrical network, we further computed an *effective* conductance score between all protein pairs. In circuit theory, effective conductance represents the ease with which current flows between two nodes when multiple parallel or series paths exist^10^. Applied to a PPI network, this measure captures the relational proximity of two proteins not only when they are directly observed to interact, but also when they are indirectly linked through multiple, independent neighbors. Each connecting path contributes to the overall effective conductance, providing a single continuous measure of protein relatedness across the entire interactome (Figure 1a-b).

**Figure 1.**
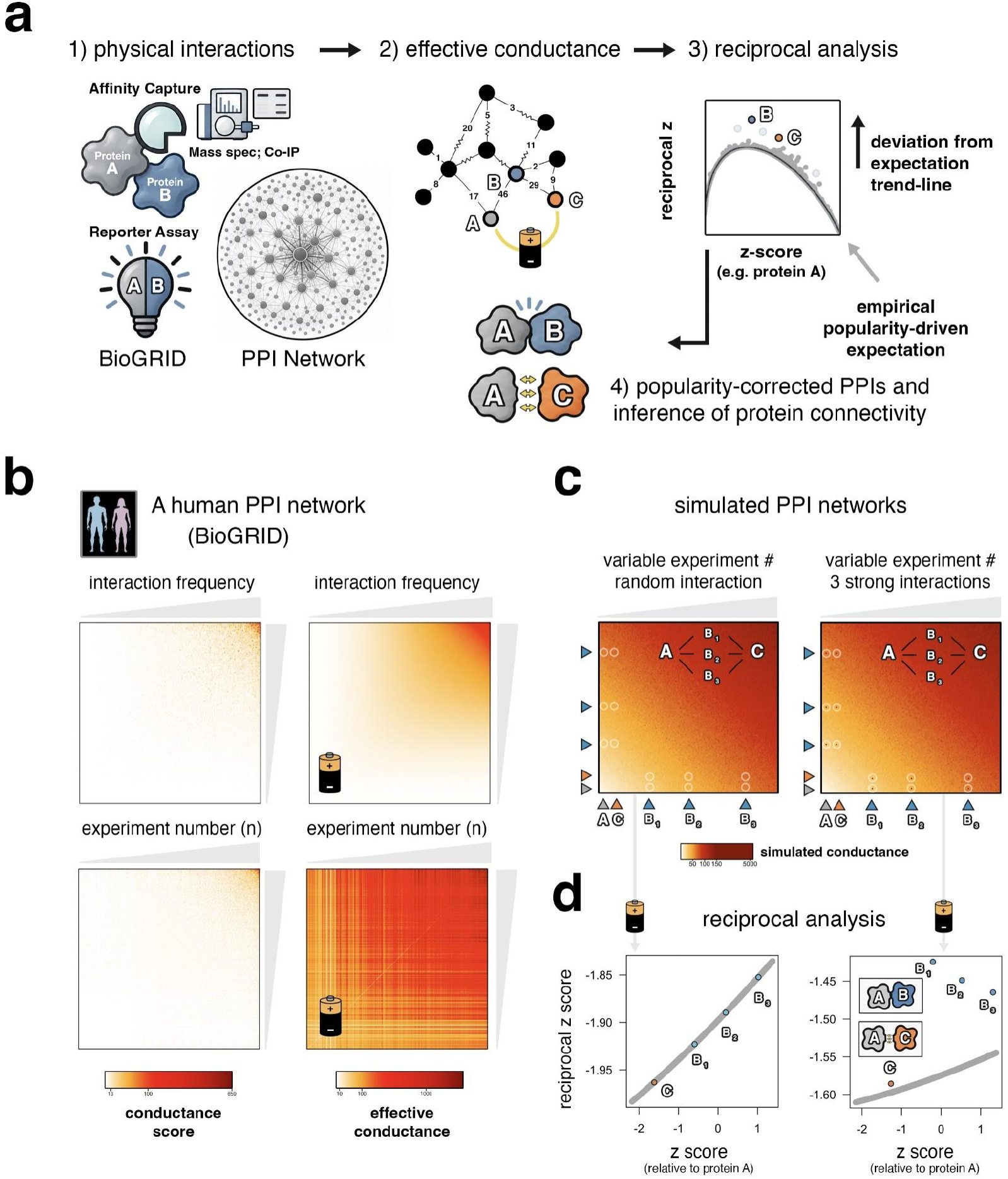
Effective conductance-based inference of protein-protein interactions and relationships. **(a)** Conceptual overview: all experimentally derived protein interactions are retrieved from BioGRID, establishing a network-based topology of the human interactome. An effective conductance score, drawn from the PPI network, is computed for all protein-protein pairs. For each protein, effective conductance scores are standardized and a reciprocal z-score analysis uncovers deviations from empirical null distributions. PPIs are thereafter inferred on the basis of significant deviations shared across both underlying proteins. **(b)** Heatmaps of conductance (left) and effective conductance scores (right), ranked by individual protein connectivity (top) and total number of experimental observations (bottom). **(c)** Heatmaps of conductance scores for simulated protein networks with no significant interactions (i.e. conductance is entirely popularity driven; left) and for simulated protein networks where proteins A and C share significant interactions with intermediate proteins B1, B2, and B3 (right). **(d)** Reciprocal z-score analysis on simulated data in panel c; simulated interactions between A::B1, A::B2, and A::B3 are recovered (right); the significant deviation of protein C uncovers a statistically significant proximity relationship between A::C in the interactome network, despite the absence of direct interactions (right).

By quantifying the connectivity between every node pair, effective conductance facilitates a global analysis of protein-protein relationships that is not possible in the initial PPI network. For example, we next considered the effective conductance between each protein-node and all other protein-nodes and thereafter computed a protein-centered z-score statistic, which represents the deviation of each protein-protein effective conductance-score relative to its own global effective conductance distribution (see ‘reciprocal plot’ methods section). Using this approach, we thereafter filter for statistically significant protein-protein connections and retain only those PPIs that are significant from the perspective of both protein-protein pairs. In addition to potentially removing spurious non-significant interactions, this framework allows for assessment of significant connectivity in the absence of prior PPI evidence. We demonstrate this principle through simulation of a fully random interaction network, where interactions are driven entirely by popularity (Supplementary Video 1), and by repeating this simulation with six hypothetical interactions linking imaginary factors B_1_, B_2_, and B_3_, to factors A and C (Figure 1c). In this scenario, these artificial interactions are recovered via reciprocal analysis, wherein A-B_1_, A-B_2_, and A-B_3_ scores deviate significantly from an empirical null expectation that emerges specifically from a popularity-biased network (Figure 1d; Supplementary Videos 1,2). Importantly, this approach additionally captures significant reciprocal deviation between factors A and C, which is driven primarily by strong effective conductance scores via connectivity with B_1_, B_2_, and B_3_ (i.e., indirect paths accounted for by effective conductance; Figure 1d).

Applying this same framework to the ~1.2 million interaction observations retrieved from BioGRID^11,13^, we recover a total of 658,913 popularity-corrected PPIs and gain 120,740 novel protein-protein relationships with significant reciprocal deviation from expectation (Figure 2a). As a first-level sanity check that our framework functions as expected, we considered the observed overlap in gene ontology between protein-protein pairs that either lost PPI status due to popularity correction, or between pairs that were identified to have strong connectivity within the popularity-corrected network in the absence of currently established PPIs (i.e. “gained protein-protein network relationships”). These analyses show that whereas gained relationships are highly significant with respect to sharing top-ranked functional GO terms^14,15^, including molecular function (MF), biological process (BP), and cellular component (CC), this pattern is not observed for protein pairs with interactions that were pruned by conductance-based reciprocal association analysis, suggesting that this framework effectively removes PPIs that are functionally unrelated and therefore more likely to be spurious (Figure 2b).

**Figure 2.**
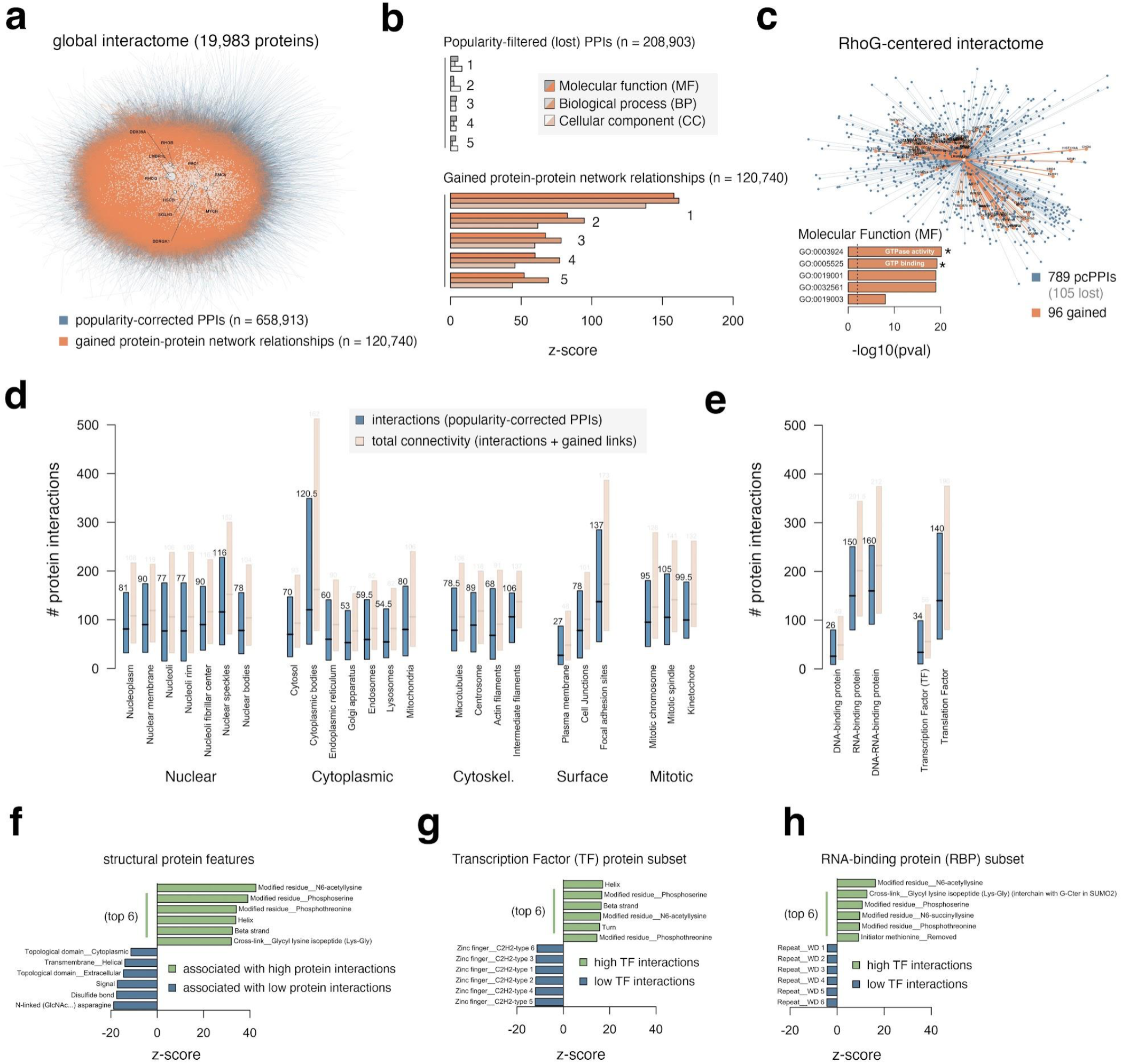
Global topology of popularity-corrected human protein interactions and significant protein-protein connectivities. **(a)** Network-based visualization of popularity-corrected protein-protein interactions (pcPPIs) following effective-conductance-based reciprocal association analysis. Significant pcPPIs are colored in blue, whereas significant protein-protein connectivity in the network, absent evidence of direct interaction, are orange. Nodes corresponding to proteins with more than 100 PPIs are highlighted in white. **(b)** Statistical overlap test between the top gene ontology terms enriched among protein interactions that are lost (grey) or with protein connections that are discovered by reciprocal analyses (orange), with the primary protein (i.e. node). The top (1), second (2), third (3), fourth (4), and fifth (5) terms, ranked by enrichment, were individually tested for shared overlap with the primary protein against an empirical null distribution. **(c)** Example starburst plot depicting interactions (blue) and novel connections (orange) for primary protein RhoG. **(d)** Distributions of PPI numbers (blue) and global connections (both PPIs + gained connections, orange) among proteins functionally linked to specific subcellular compartments. **(e)** Analogous distributions for DNA-, RNA-, and DNA-RNA-binding proteins, and related transcription factor (TF) and translation factor GO terms. **(f-h)** Top protein-intrinsic structural features associated with either high (green) or low (blue) protein connectivity among all proteins (panel f), or among transcription factor (g) or RNA-binding protein (h) subsets, specifically.

### Emergent properties of a popularity-corrected human protein interaction network

Building on the reciprocal-analysis-informed topology of PPIs and protein connectivities, we next investigated basic properties that emerge from our framework, such as the identities of highly-connected “hub” proteins. Among a list of such factors that include LMBR1L, MYCN, DDX39A, and HSCB (Figure 2a), we find that RhoG – a member of the Rho family of GTPases^16^ – is the single most connected protein in our network (789 total popularity-corrected PPIs, 96 novel protein-protein connections, Figure 2c). More broadly, we find that proteins associated with nuclear speckles (i.e. splicing bodies) exhibit the highest level of connectivity in the nucleus (median 116 pcPPIs), whereas factors generally associated with cytoplasmic bodies are similarly highly connected in the cytoplasm (median 120.5 pcPPIs, Figure 2d)^17^. Likewise, RNA-binding proteins (RBPs), which are abundant in both nuclear speckles and cytoplasmic bodies^18^, exhibit high connectivity (median 150 pcPPIs), in contrast to DNA-binding proteins and transcription factors (TFs; median 26 and 34 pcPPIs, respectively; Figure 2e). Analysis of protein-intrinsic structural features associated with high-connectivity finds that phosphoserine, phosphothreonine, and N6-acetylation modifications are universally associated with higher-than-expected connectivity, whether considering all proteins together or TFs and RBPs as individual subsets (Figure 2f-h). Conversely, TFs with C2H2-type zinc finger motifs and RBPs with WD-repeat motifs were associated with lower-than-expected connectivity patterns within these two sub-categories (Figure 2g-h).

### Popularity-corrected protein interactions associated with DNA-dependent RNA polymerase enzymes

To capitalize on the utility of the global pcPPI network, we next explored protein-protein interactions related to our primary molecular research interest, RNA polymerase (Pol) III. In brief, eukaryotic transcription generally involves three distinct enzymes - Pol I, Pol II, and Pol III - which are structurally and functionally distinguished by a multitude of enzyme-specific subunits and unique gene targets^19-21^. Whereas Pol I and Pol II produce ribosomal rRNA and messenger mRNA, respectively, both Pol II and Pol III produce a plethora of small, highly structured noncoding (nc)RNAs^22-25^. Here, we sought to explore pcPPIs for Pol III, as well as for Pols I and II, by focusing on enzyme-specific factors, such as core subunits POLR1B (Pol I-specific), POLR2B (Pol II-specific), and POLR3A (Pol III-specific), which yield 105, 204, and 105 pcPPIs, respectively (Figure 3a-c). Interestingly, inspection of each enzyme-specific subunit-interactome revealed a surprising number of cross-enzyme pcPPIs, such as between POLR1B (Pol I) and Pol II subunits POLR2A, -2B, -2C, -2D, -2G, -2I, -2J, and -2M (Figure 3d), and, vice-versa, between POLR2B (Pol II) and Pol I subunits POLR1A, -1B, -1E, -1G (CD3EAP), and -1H (ZNRD1; Figure 3e). Analogous cross-enzyme interactions are also observed for Pol III, such as between large subunit POLR3A and Pol II subunits POLR2A, -2B, -2C, -2D, -2G, - 2I, -2J, and -2M, suggesting that these pan-complex protein-protein interactions are potentially common across all three enzymes (Figure 3f).

**Figure 3.**
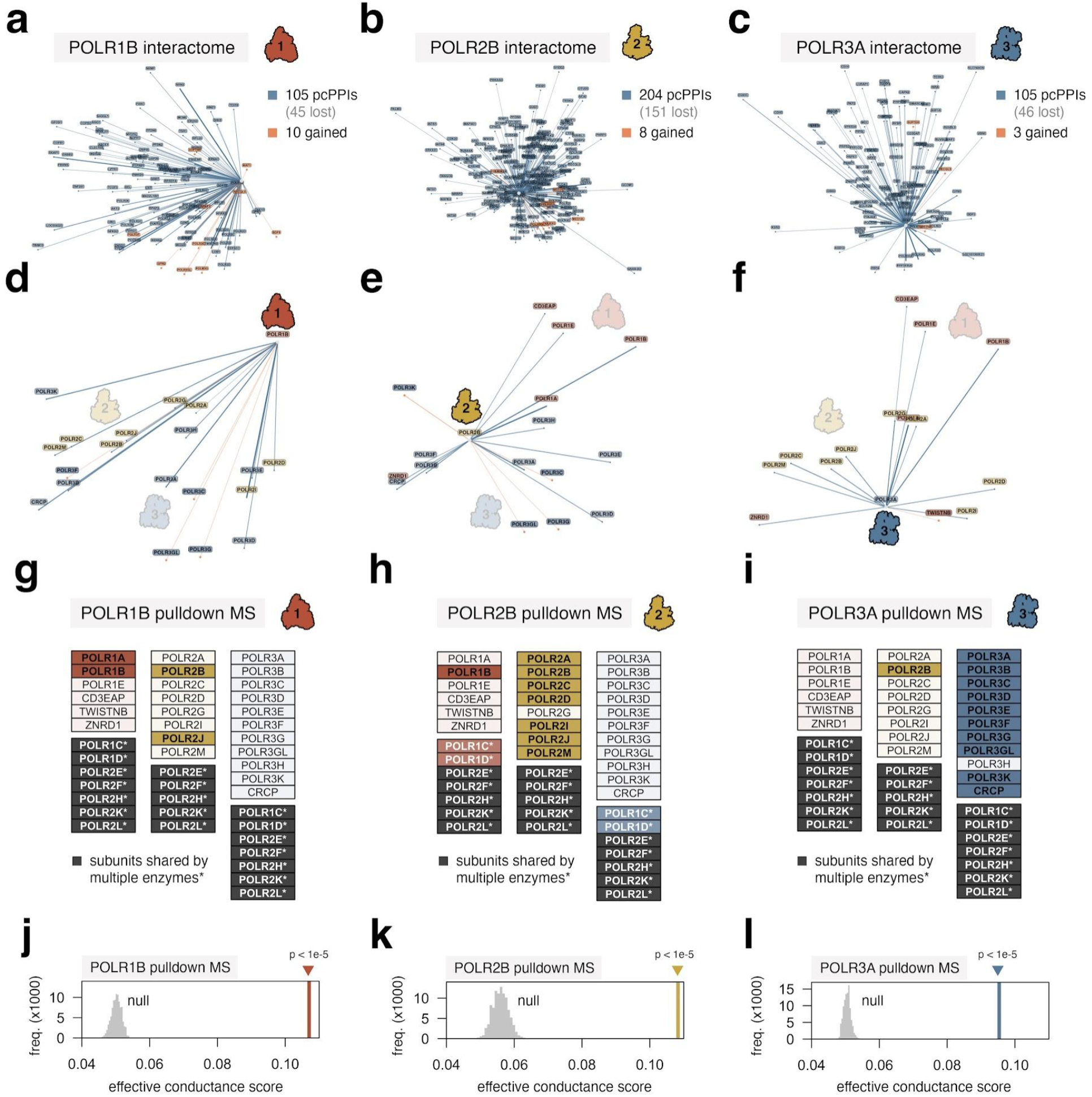
Popularity-corrected protein-protein interactions reveal unexpected connectivity between enzyme-specific RNA polymerase subunits. **(a-c)** Starburst plots depicting popularity-corrected protein-protein interactions (pcPPIs; blue) and novel connections (orange) for Pol I subunit POLR1B (a), Pol II subunit POLR2B (b), and Pol III subunit POLR3A (c). **(d-f)** Analogous starburst plots, restricted to distinct-enzyme-specific subunits for POLR1B (d), POLR2B (e), and POLR3A (f). Subunit IDs are colored by enzyme (Pol I = red; Pol II = gold; Pol III = blue). **(g-i)** Visual illustration of enzyme-specific and shared subunits significantly enriched in pull-down mass-spec (MS) experiments for POLR1B (g), POLR2B (h), or POLR3A (i) in HEK293T cells. Darkened colors denote confirmed hit; Note that among shared subunits, POLR1C and POLR1D are canonically Pol I and Pol III-specific. **(j-l)** Empirical null distributions and observed effective conductance scores for proteins identified by pulldown MS, for either POLR1B (j), POLR2B (k), or POLR3A (l). Network conductance scores represent the median effective conductance among all proteins for which a given pull-down MS PPI hit was not previously reported to BioGRID.

In light of these findings, we independently carried out pull-down mass-spectrometry (pull-down MS) experiments against POLR1B, POLR2B, and POLR3A in HEK293T cells. Though sparse, our pull-down MS results do recover enrichment for distinct enzyme-specific subunits^26^, including between POLR1B-POLR2B and POLR1B-POLR2J (Figure 3g), and between POLR2B-POLR1B, POLR2B-POLR1C, POLR2B-POLR1D (Figure 3h). Likewise, affinity-capture of POLR3A recovers POLR2B enrichment (Figure 3i), altogether suggesting a potentially unappreciated macro-molecular interplay between subunits for all three enzymes. Beyond providing supporting evidence for cross-enzyme connectivities, we additionally show that previously unreported protein-protein interactions identified by pull-down MS are characterized by significantly higher effective conductance scores than expected, both for our Pol III subunits of interest (Figure 3j-l) as well as other unrelated proteins (Supplementary Figure 1). These findings highlight the information-rich nature of effective-conductance-based network analysis and its potential value in predicting a broad range of macromolecular interactions in humans.

### Leveraging reciprocal effective conductance scores to predict macromolecular interactions

The interactions demonstrated for POLR1B, POLR2B, and POLR3A represent only a small fraction of cross-enzyme pcPPIs observed between Pols I, II, and III, as all three complexes have many additional enzyme-specific subunits. Indeed, taking all such subunits into account reveals a large number of complex-complex connections, including both pcPPIs (Figure 4a) as well as gained protein-protein associations that were quantitatively driven by underlying network PPI paths (Figure 4b). Taking these patterns into account, we reasoned that a macromolecular complex-complex interaction (CCI) score, computed on the basis of all underlying protein-protein connectivities determined by reciprocal effective conductance analysis, may reveal insight on these and potentially other macromolecular interactions (Figure 4c). Computing CCI scores between Pol I and > 5,000 other unique mammalian complexes, we find that both Pol II and Pol III do emerge among the highest multi-subunit complexes with significant Pol I-CCI scores (Figure 4d). These relationships, which are determined on the strict basis of enzyme-specific subunits (shared subunits are omitted from CCI calculations), are also observed from the perspective of either Pol II or Pol III (Figure 4e and Figure 4f, respectively).

**Figure 4.**
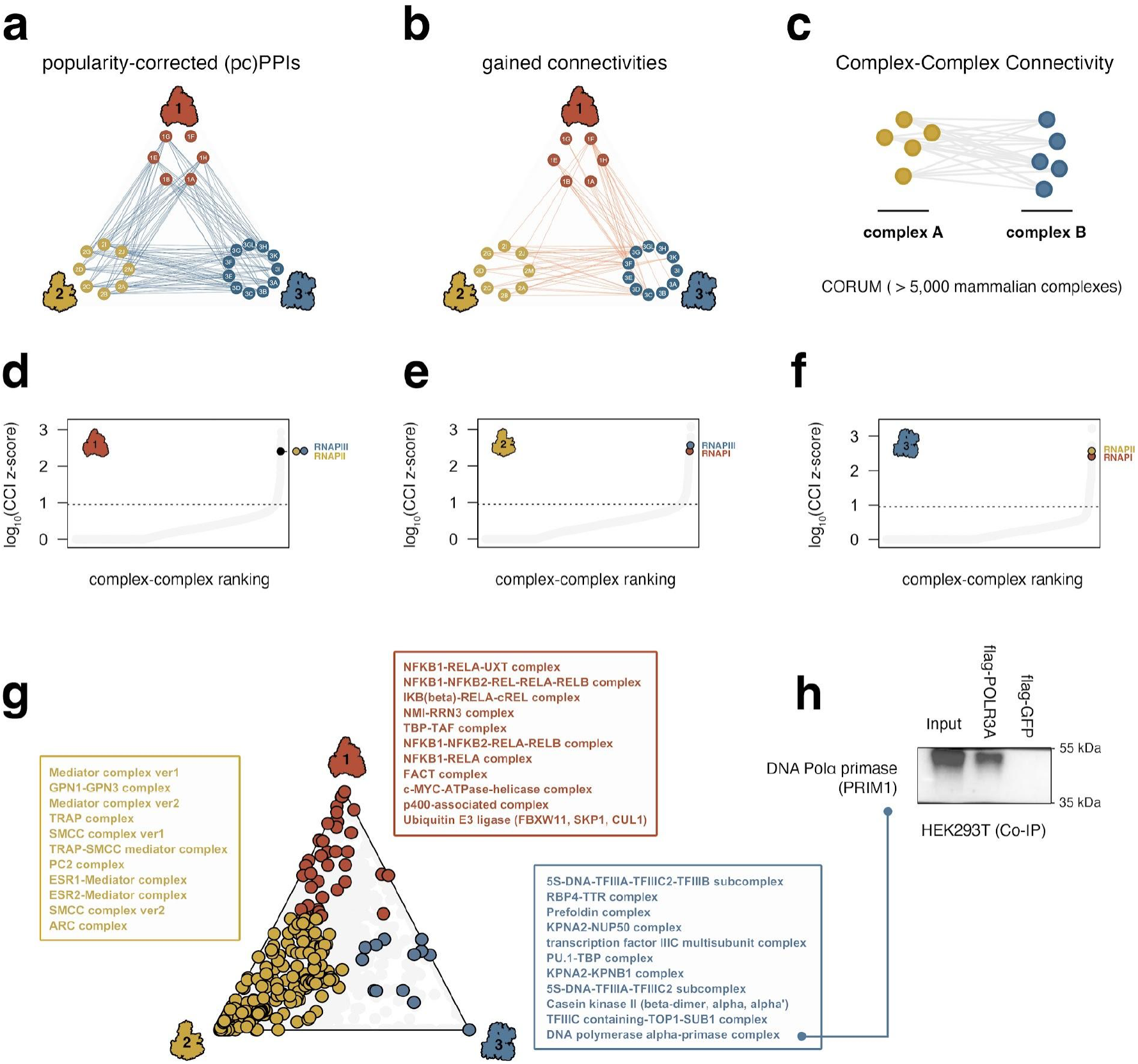
Statistical analysis of complex-complex connectivities established via protein-protein reciprocal effective conductance scores reveals unexpected macromolecular interactions. **(a)** Network-based visualization of popularity-corrected protein-protein interactions (pcPPIs) between all enzyme-specific subunits of RNA polymerase I, II, and III. Edges (blue) denote significant pcPPIs observed between a given subunit-subunit pair (nodes). **(b)** Network-based visualization of statistically significant “gained” connectivities. Edges (orange) denote significant connectivity - in the absence of direct protein-protein interaction evidence - between a given subunit-subunit pair (nodes). **(c)** Illustration of Complex-Complex Interaction (CCI) framework, which compares cross-complex subunit-subunit connectivities against an empirical null expectation. Complex annotations were retrieved from CORUM. **(d-f)** CCI scores corresponding to ~5,000 mammalian complexes, ranked by log10(z-score), for RNA Pol I (d), RNA Pol II (e), and RNA Pol III (f). Distinct RNA polymerase enzymes are highlighted in each respective plot. **(g)** Dominance-based visualization of significant macromolecular interactions. Data points denote unique mammalian complex, colored by the respective enzyme with strongest complex-complex connectivity. Insets describe the most significant Pol I-dominated connectivities (red), Pol II-dominated connectivities (gold), and Pol III-dominated connectivities (blue). **(h)** Co-immunoprecipitation (co-IP) analysis of DNA pola primase (subunit PRIM1) following pull-down of large Pol III subunit POLR3A in HEK293T cells.

While these data add additional compelling evidence in support of putative cross-enzyme interactions, we next leveraged our framework to identify other, potentially novel macromolecular interactions involving Pol III. To prioritize interactions unique to Pol III, we simultaneously profiled multi-protein complexes with significant CCI scores to any or all RNA polymerase machineries, and thereafter focused on factors with particularly strong, Pol III-specific connectivities. Importantly, this analysis detects well-established macromolecular relationships, such as between Pol I and the NMI-RRN3 complex^27,28^, between Pol II and the Mediator complex^29^, and between Pol III and Pol III-specific general transcription factor complexes TFIIIB and TFIIIC^30^ (Figure 4g). Other enzyme-dominated complex-complex interactions that are congruent with previous reports include those between Pol I and the FACT complex^31^, between Pol II and the GPN1/3 complex^32^, and between Pol III and the Casein kinase II complex^33-36^. However, we also identify multiple Pol III-centered connections that, to our knowledge, have not been previously described, such as putative interactions with the RBP4-TTR complex – involveed in retinol transport^37^ – and the DNA polymerase alpha-primase complex – involved in DNA replication initiation^38,39^ (Figure 4g). We further show that PRIM1, the catalytic subunit of DNA primase, does indeed interact with POLR3A via co-IP (Figure 4g), altogether suggesting a potential for unexpected crosstalk between Pol III transcription and DNA replication.

### CoBRAA: A conductance-based reciprocal association analysis tool for PPI and CCI exploration and prediction

The identification of putative complex-complex interactions between Pol III and DNA primase exemplifies the utility of leveraging protein-protein connectivity to predict macromolecular properties. To maximize the accessibility of this framework to researchers of all interests and specializations, we developed CoBRAA (for conductance-based reciprocal association analysis) – an interactive tool with a graphical user interface to facilitate exploration of protein-centered reciprocal effective conductance scores and the resulting protein interactions that pass statistical filtering (i.e. pcPPIs; Figure 5a-b), as well as higher-level complex-complex interaction scores identified for ~5,000 CORUM-annotated mammalian complexes (Figure 5c)^40,41^. While these data can be used to extrapolate all pcPPIs between 2- or more multi-subunit complexes (Figure 5d), CoBRAA also includes advanced options for visualizing complex-centered interactions among lists of user-defined macro-molecules of interest (Figure 5e). As an example, we demonstrate that the NuRD (Nucleosome Remodeling and Deacetylase) complex subunits interact with both DNMT3B and Polycomb repressive complex 2/3 (PRC2/3)-specific sub-units (Figure 5d). While all three are well-established regulatory complexes associated with gene repression^42-45^, feature-dominance-analysis captures NuRD-, DNMT3B-, and PRC2/3-specific connections, such as those between NuRD and Casein kinase II, between DNMT3B and condensin II, and between PRC2/3 and the LCOR/VDR co-repressor complex (Figure 5e-f)^46^. As shown for Pol III and DNA primase interactions, these and other CoBRAA-derived observations may inspire deeper study and novel insights.

**Figure 5.**
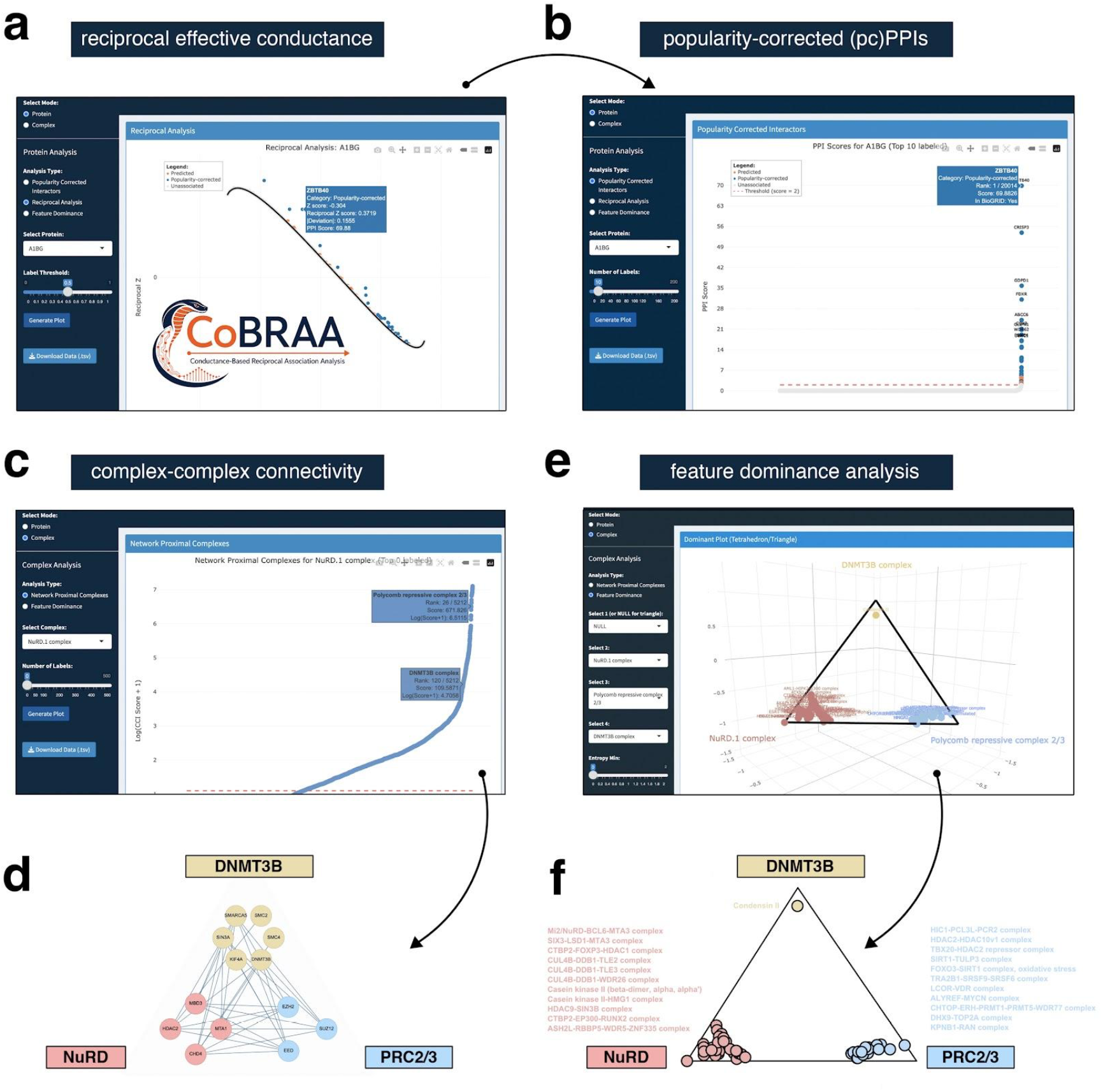
CoBRAA (Conductance Based Reciprocal Association Analysis): An interactive tool for protein-protein and complex-complex interaction and connectivity exploration. **(a-b)** Example exploration of reciprocal effective conductance analysis (panel a) and the resulting popul-*(continued from previous page)*: -arity-corrected protein-protein interactions (pcPPIs, panel b) for A1BG - a plasma glycoprotein - using the CoBRAA graphical user interface. **(c)** Example exploration of complex-complex interaction (CCI) scores for NuRD - the Nucleosome Remodeling and Deacetylase complex - using CoBRAA. **(d)** Network-based visualization of popularity-corrected protein-protein interactions between all complex-specific subunits of NuRD, DNMT3B, and PRC2/3. Edges (blue) denote significant pcPPIs observed between a given subunit-subunit pair (nodes). Example exploration of complex-level feature dominance analysis for NuRD, DNMT3B, and PRC2/3 multi-subunit complexes. **(f)** Dominance-based visualization of significant macromolecular interactions. Data points denote a specific, unique mammalian complex, colored by the respective complex (NuRD, DNMT3B, or PRC2/3) with strongest complex-complex connectivity. Insets describe the most significant NuRD-dominated connectivities (pink), DNMT3B-dominated connectivities (yellow), and PRC2/3-dominated connectivities (blue).

## Discussion

Here, we establish a reciprocal effective conductance-based framework that, when applied to ~1.2 million reported observations on BioGRID, identifies a total of 658,913 significant human protein-protein interactions. While this analysis helps to adjust for the uneven distribution of protein-centered interaction studies performed to date, a major advantage of computing *effective* conductance lies in its representation of theoretical conductance through all node pairs within the network. This measure of inferred connectivity among all protein pairs is a key element that facilitates higher-level macromolecular analyses, which consider the observed connectivities among each of the complex-constituent subunits.

Application of our framework uncovers both emergent properties of the human interactome, such as the high degree of connectivity among RNA-binding proteins and nuclear speckle-associated factors (for which multivalent properties are already well characterized), and of userdefined, protein-centered interaction properties that may not be presently well understood. To the latter point, we demonstrate higher-than-expected interactions among enzyme-specific subunits of all three eukaryotic DNA-dependent RNA polymerases, as well as macromolecular interactions between RNA polymerase III and the DNA primase complex. These findings raise interesting possibilities of direct transcription-replication coordination at sites transcribed by Pol III - including tRNA genes - that will inspire deeper investigation.

Finally, we present CoBRAA as an accessible tool for exploring these and other protein-protein and complex-complex relationships derived from reciprocal association analysis. Given that protein interactions coordinate nearly every cellular process, we expect this resource to facilitate deeper investigation and hypothesis generation across a broad range of proteins, pathways, and macromolecular systems.

### Limitations of this study

Our framework is determined by the underlying BioGRID interaction matrix and therefore cannot overcome absent or extremely sparse experimental coverage for specific factors. Here, interaction frequencies simply reflect the number of independent protein-protein observations across multiple experimental approaches and conditions, and thus variables such as biochemical affinity and context-specificity are not considered. Though powerful, our reciprocal effective conductance analysis further prioritizes observations that are statistically significant from the perspective of each independent factor, and thus biologically meaningful interactions involving highly connected factors (e.g., those that truly interact with a majority of the proteome, such as molecular chaperones) may be lost in the process. While we annotate significant reciprocal associations that are novel (i.e. not supported by previous PPI studies), these “gained associations” cannot be interpreted as evidence of direct physical interaction. With these limitations in mind, all protein-protein and complex-complex interactions and connections predicated on reciprocal effective conductance naturally require additional biochemical validation and functional analysis.

## Methods

### Construction of PPI matrix using BIOGRID

The PPI matrix was built using the BIOGRID (v4.4.243) database^11^. Only the physical interactions were retained, specifically including Affinity Capture-Luminescence, (2) Affinity Capture-MS, (3) Affinity Capture-RNA, (4) Affinity Capture-Western, (5) Biochemical Activity, (6) Co-crystal Structure, (7) Co-fractionation, (8) Co-localization, (9) Co-purification, (10) Cross-Linking-MS (XL-MS), (11) Far Western, (12) FRET, (13) PCA, (14) Protein-peptide; Protein-RNA, (15) Proximity Label-MS, (16) Reconstituted Complex and (17) Two-hybrid. Disconnected subnetworks, if present, were removed, retaining only the largest connected PPI network. Self-interactions were removed if present. The resulting protein-protein interaction (PPI) network is represented as a weighted interaction matrix, *C* where each entry corresponds *C*_*i,j*_ to the number of observed interactions between proteins *i* and *j* in the BIOGRID database. The PPI matrix dimension is 20,014 × 20,014, representing interactions between 20,014 proteins (nodes).

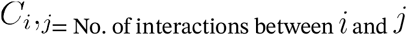

### Construction of PPI matrix in simulation 1

In the first simulation, we construct a hypothetical interaction network consisting of 1000 proteins, labeled *P*_*1*_,*P*_2_,…,*P*_*n*_,…,*P*_1000_. Each protein *P*_*i*_ is assigned a popularity score equal to its index *i*, meaning *P*_*i*_ protein is assumed to form *i* interactions with every other protein. To model this, we define an initial interaction matrix *C*_*p*_, where each row is filled with the protein’s popularity score i.e.,. (*C*_*p*_)_*ij=*_*i*.

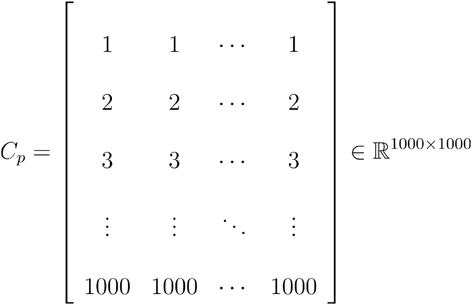

This implies that each protein is interacting equally with all others, and there are no specific interaction preferences in this preliminary matrix. To make the interaction symmetric (i.e., bidirectional interactions), we define a final matrix as the sum of the matrix and its transpose:

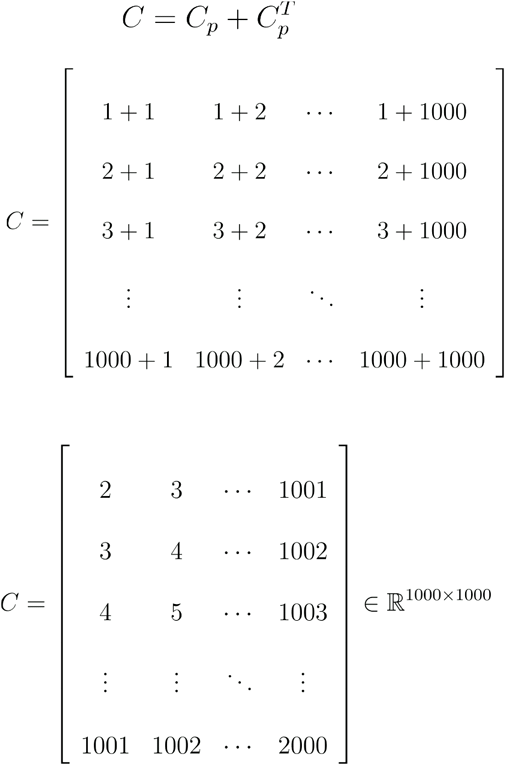

This results in a symmetric interaction matrix *C*, where the interaction score between proteins *P*_*i*_ and *P*_*j*_ is simply the sum of their popularity scores, i.e.,. *C*_*ij*_*=i+j*.

### Construction of PPI matrix in simulation 2

In the second simulation, we constructed a similar hypothetical interaction network consisting of 1000 proteins, labeled *P*_*1*_,*P*_2_,…,*P*_*n*_,…,*P*_1000_. Each protein *P*_*i*_ was assigned a popularity score equal to its index. In simulation 1, *P*_*i*_ protein was assumed to form interactions with every other protein, whereas here we assume that protein *P*_*i*_ was profiled (experimented as a bait) *i* times and in every experiment the protein shows interaction with 50 randomly sampled proteins. The assumption was that all the proteins have the same number of interactors (50 in this case) and thus identify the same number of interactors in any profiling experiment. To model this, we define an initial interaction matrix *Cp*, and fill this matrix in a row-wise order with interactions generated from random sampling, for example, when we fill the rows for protein *P*_7_, we profile this protein times and in every profiling we add it’s interactions with 50 randomly sampled proteins, thus protein *P*_7_ will have 7 * 50 interactions.

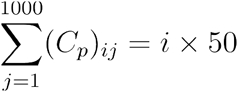

This implies that each protein has a tendency to interact with all other proteins equally, and hence the tendency is to have no specific interaction preferences in this preliminary matrix, however the random sampling will generate some level of noise and thus the proteins don’t have equal interactions with other proteins. Finally similar to the simulation 1, to make the interaction symmetric (i.e., bidirectional interactions), we define a final matrix *C* as the sum of the matrix and its transpose:

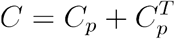

### Construction of Effective Conductance matrix from PPI matrix

We assumed PPI network as a purely resistive electrical network where proteins are nodes and the interaction frequency between any protein is electrical conductance between them. Thus, using PPI matrix as conductance matrix (*C*), we created effective conductance matrix (*EC*) whose elements are the pairwise effective conductance between the proteins.

First, we define the Laplacian matrix *L*∈ ℝ^*n⨯n*^ as:

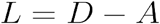

Where *D* is the degree matrix, a diagonal matrix where *D*_*ii*_ degree of protein, and all off-diagonal elements are zero:

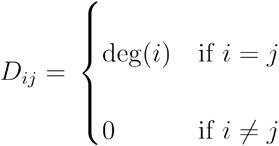

*A* is the is the adjacency matrix, constructed from the original interaction matrix *C* by setting its diagonal elements to zero:

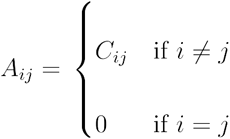

Second, the Moore-Penrose pseudoinverse of the Laplacian matrix (*L*), denoted as *L*+, was computed using the generalized inverse function ginv() from the MASS package in R.

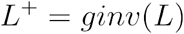

Finally, the effective conductance matrix (*EC*) was computed using the following formula^47^:

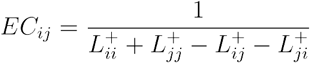

**Reciprocal plot**

Here, we create Reciprocal plot using effective conductance matrix (*EC*):

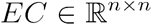

Effective conductance to self is infinite, which we first converted to zero:

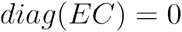

Then, we row-wise Z scaled the effective conductance matrix (*EC*) to obtain matrix *Z* such that:

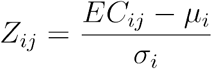

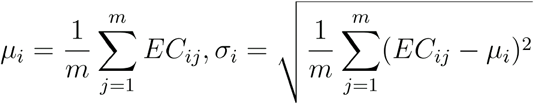

Finally, Reciprocal plot for any protein *P*_*i*_ is plotted as a scatter plot by taking the *i*^*th*^ row of *Z* as the x-axis and the *i*^*th*^ column of *Z* as the y-axis of the plot.

### Residual calculation from spline fits in reciprocal plots

For each reciprocal scatter plot, we fit a smoothing spline. Spline fitting is performed using the smooth.spline() function from the stats package in R, using the maximum allowable degrees of freedom. Spline fitting is carried out in two steps. First, a smoothing spline is fit using all points in the reciprocal plot. Vertical residuals from this spline are then computed for each point, and points with large deviations from the fitted spline are excluded. A second smoothing spline is then fit using only the remaining points. Now, the residuals are calculated for each point from the second spline-fitted line and z-normalized.

### Calculation of pairwise closeness score

For a given pair of proteins *a* and *b*, we quantify their pairwise closeness using deviations measured in each other’s reciprocal plots. First, we obtain the Z-normalized residual of protein *a* from the spline-fitted line in the reciprocal plot of protein *b*, denoted as *R*_*a,b*_. Similarly, we obtain the Z-normalized residual of protein *b* from the reciprocal plot of protein *a*, denoted as *R*_*b,c*_. The pairwise closeness score between proteins and is then defined as:

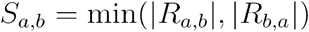

### Statistical overlap test of GO for protein lost and gained partners

GO enrichment was performed on the gained interactors of each protein using enrichGO() function from the clusterProfiler R package^48^. We then determined whether the first enriched GO term among the gained interactors also matched as a GO term of the focal protein and counted the number of such matches across all proteins. A null distribution was generated by randomly shuffling the focal-protein GO annotations across proteins 10,000 times while keeping the enriched GO terms of the gained interactors unchanged. An empirical Z-score was calculated by comparing the observed number of matches with the permutation distribution. The same analysis was repeated for the second through fifth enriched GO terms, for Cellular Component (CC) and Molecular Function (MF), and separately for lost interactors as well.

### Visualization of popularity corrected PPI network

For visualization of the popularity-corrected PPI network, an undirected graph was constructed using the igraph R package^49-51^, and proteins with zero degree were removed. The coordinates of the remaining proteins were generated using the Large Graph Layout (LGL) algorithm implemented with layout_with_lgl(). The network was visualized using ggraph (https://cran.rstudio.com/web/packages/ggraph/), with popularity-corrected and newly gained relationship edges shown in different colors and plotted with low width and transparency to improve visualization of the dense network. For clarity, only proteins with degree ≥100 were displayed as visible node points, with node size scaled according to degree. The 10 proteins with the highest degree were further highlighted and labeled.

### Association of protein structural features with PPI connectivity

Protein structural and sequence features were retrieved from UniProtKB through the UniProt REST API^52,53^. HGNC gene symbols were mapped to UniProt/Swiss-Prot accessions using Ensembl BioMart through the biomaRt R package prior to feature retrieval^54,55^. UniProt annotations including secondary-structure elements, modified residues, cross-links, topological domains, transmembrane regions, repeats, and other annotated features were extracted, while sequence conflicts, natural variants, chains, sites, mutagenesis annotations, and alternative sequences were excluded. To identify protein features associated with PPI connectivity, the mean interaction degree of proteins containing each feature was calculated. For a feature present in n proteins, a null distribution was generated from 100,000 random samples of n proteins from the corresponding background population. Z-scores were calculated by comparing the observed mean degree with the mean and standard deviation of the permutation distribution; positive and negative Z-scores indicate association with higher and lower PPI connectivity, respectively. This analysis was first performed across all proteins and subsequently repeated within transcription factor and RNA-binding protein subsets. Transcription factors were defined using the human TF list from the Aerts lab’s cisTarget resource (https://resources.aertslab.org/cistarget/tf_lists/allTFs_hg38.txt)^56-58^, and RNA-binding proteins were defined using the high-confidence human RBP set from RBP2GO (https://rbp2go-2-beta.dkfz.de/)^59,60^.

### Calculation of Complex-Complex Interaction Score

Human protein complexes were obtained from CORUM (version 5.3)^40,41^. For each pair of protein complexes, we considered all possible inter-complex interactions between their constituent proteins and calculated the median of the interaction score. We then generated a background distribution for each combination of complex sizes by randomly selecting two sets of proteins of the corresponding sizes. This random sampling was repeated 10,000 times, and the median interaction score was calculated for each iteration. The mean and standard deviation of these 10,000 median values were calculated, and the observed median interaction score for each complex pair was converted to a Z-score using the corresponding background mean and standard deviation. The resulting Z-scores were used to construct the complex–complex interaction matrix.

### Cell lines and culture conditions

HEK293T cells were obtained from ATCC (CRL-3216, Batch#70049877) and grown in 10 cm BioLite™ Cell Culture Treated Dishes (Catalog# 12-556-002 Thermo Scientific) in Dulbecco’s Modified Eagle Medium, high glucose (Catalog# 11965092, Gibco). HEK293T experiments were conducted on cells between passage 10-25.

### Pull-down Mass Spectrometry and Immunoblotting

Flag-tagged POLR1B, POLR2B, and POLR3A pull-down MS experiments were recently performed as previously in Cheng et al., 2025^26^. In addition to Pol I, II, and III subunits, Flag-tagged ATRN (Clone ID: OHu18959D, GenScript), Flag-tagged FBRSL1 (Clone ID: OHu07567D, GenScript), Flag-tagged TTC13 (Clone ID: OHu04088D, GenScript), and Flag-tagged eGFP control were ectopically expressed in HEK293T cells via transfection using Lipofectamine 3000 Transfection Reagent (Cat #L3000015) following manufacturer’s protocol. Cells are collected 2 days after transfection and coIP performed using Pierce™ Classic Magnetic IP/Co-IP Kit(88804) and Pierce™ Anti-DYKDDDDK Magnetic Agarose(A36797) following manufacturer’s protocol. Elution and input are incubated at 95°C with diluted 4× Laemmli Sample Buffer (Catalog#1610741, BIO-RAD Hercules, CA, USA) for 5min. Mass spectrometry collection and analysis is performed as described^26^. For immunoblotting, proteins were separated on 4-20% Mini-PROTEAN® TGX Stain-Free™ Protein Gels, 15 wells (Catalog# 4568096, BIO-RAD, Hercules, CA, USA) using 10× Tris/Glycine/SDS (Catalog# 1610732, BIO-RAD, Hercules, CA, USA) and transferred onto polyvinylidene difluoride membranes (0.2 um) (Catalog# LC2002, Invitrogen, Waltham, MA, USA) with a Trans-Blot® Turbo™ Transfer System (Catalog# 1704150, BIO-RAD, Hercules, CA, USA). Transfer membranes were blocked with a 5% blotting-grade blocker (Catalog# 1706404, BIORAD, Hercules, CA, USA), followed by incubation with the PRIM1 Polyclonal Antibody (Catalog# 10773-1-ap, Rosemont, Proteintech, IL, USA) at 4 °C overnight. Membranes were washed with TBST and incubated with rabbit secondary antibodies conjugated with horseradish peroxidase (Catalog# 31462, Invitrogen, Waltham, MA, USA) for 2 h at room temperature followed by three washes in TBST. Proteins were visualized using either SuperSignal West Pico (Catalog# 34580, Thermo Scientific, Waltham, MA, USA) or SuperSignal West Femto (Catalog# 34096, Thermo Scientific, Waltham, MA, USA) with a ChemiDoc™ Touch Imaging System (Catalog# 1708370, BIO-RAD, Hercules, CA, USA).

## Acknowledgements

We thank Drs. Peter Yau, Justine Arrington, Brian Imai, and staff at the Carver Biotechnology Center for proteomics services. We thank Drs. Alvaro Hernandez, Chris Wright, Danman Zhang, and staff at the Carver Biotechnology Center and administrators of the Carl R. Woese Institute for Genomic Biology (UIUC) Biocluster for computational support. We thank Professor Sihai Dave Zhao and members of the Van Bortle lab for helpful suggestions. This work was supported by the Roy J. Carver Charitable Trust grant 26-6113 and the National Institutes of Health grant R35GM164071 to KVB.

## Author contributions

Study design: RKC, KVB. Data collection: RKC, SZ, RC, MAP. Data analysis: RKC, SZ, RC, SL, KVB. Data interpretation: RKC, KVB. Writing: RKC and KVB with comments from all authors.

## Supplementary Figures

**Supplementary Figure 1.**
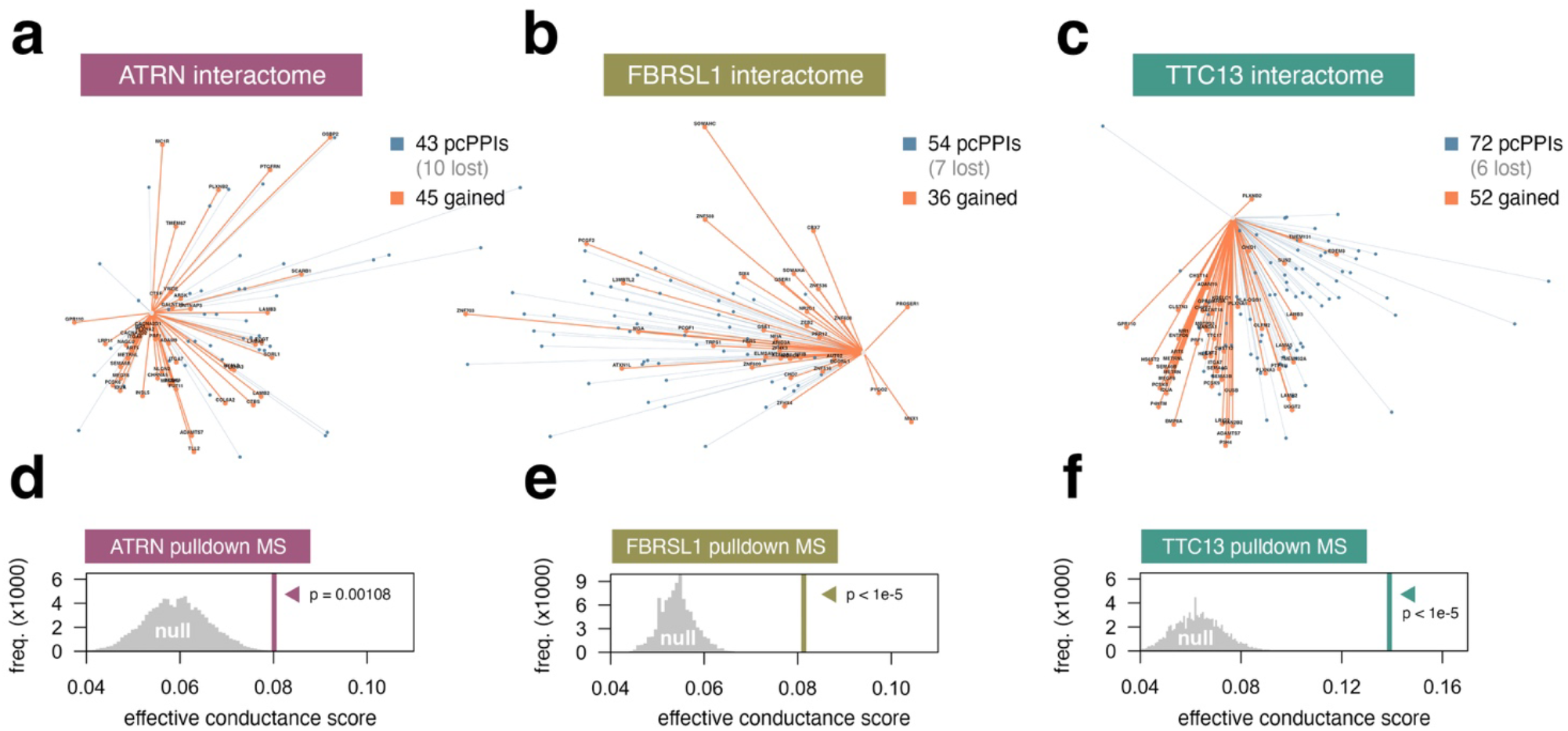
Pull-down mass spectrometry and corresponding effective conductance scores observed for interactions captured by ATRN, FBRSL1, and TTC13. **(a-c)** Starburst plots depicting popularity-corrected protein-protein interactions (pcPPIs; blue) and novel connections (orange) – derived via Conductance-Based Reciprocal Association Analysis (CoBRAA) – for proteins ATRN (a), FBRSL1 (b), and TTC13 (c). **(d-f)** Empirical null distributions and observed effective conductance scores for proteins identified by pulldown MS, for either ATRN (d), FBRSL1 (e), or TTC13 (f).

## Supplementary Videos

**Supplementary Video 1 (See SupplementaryVideo1.gif)**. Reciprocal analysis simulation of a hypothetical protein network (n=1000 proteins), applied to the z-standardized conductance scores (top) or z-standardized effective conductance scores (bottom). Moving plots trace individual proteins based on simulated popularity (i.e. number and values of protein-protein connections, from low-to-high popularity). The x-axis represents the distribution of z-standardized conductance (top) or effective conductance scores (bottom) for a primary protein, and the y-axis represents the reciprocal z-score for the same primary protein within the distribution of all other proteins. Simulation 1 (left) depicts a protein network entirely determined by popularity, in contrast to Simulation 2 (right), which models protein-protein interactions by random assignment (i.e., connectivities that are not popularity driven).

**Supplementary Video 2 (See SupplementaryVideo2.gif)**. Reciprocal analysis simulation of a hypothetical protein network (n=1000 proteins), as in Supplementary Video 1, but in the absence of popularity bias (i.e., all proteins have equivalent representation in the network). Here, simulation 1 (left) – which lacks meaningful connections – cannot be z-standardized (all primary proteins have the same connectivity total). In simulation 2 (right) – which again models protein-protein interactions by random assignment – we now observe the distribution of z-standardized conductance (top) or effective conductance scores (bottom) for a primary protein, and the reciprocal z-score for the same primary protein within the distribution of all other proteins – in a network that does not include popularity bias.

## References

1. De Las Rivas, J., and Fontanillo, C. (2012). Protein-protein interaction networks: unraveling the wiring of molecular machines within the cell. Brief Funct Genomics 11, 489–496. 10.1093/bfgp/els036.

2. Berggard, T., Linse, S., and James, P. (2007). Methods for the detection and analysis of protein-protein interactions. Proteomics 7, 2833–2842. 10.1002/pmic.200700131.

3. Hakes, L., Pinney, J.W., Robertson, D.L., and Lovell, S.C. (2008). Protein-protein interaction networks and biology--what’s the connection? Nat Biotechnol 26, 69–72. 10.1038/nbt0108-69.

4. Blumenthal, D.B., Lucchetta, M., Kleist, L., Fekete, S.P., List, M., and Schaefer, M.H. (2024). Emergence of power law distributions in protein-protein interaction networks through study bias. Elife 13. 10.7554/eLife.99951.

5. Gillis, J., Ballouz, S., and Pavlidis, P. (2014). Bias tradeoffs in the creation and analysis of protein-protein interaction networks. J Proteomics 100, 44–54. 10.1016/j.jprot.2014.01.020.

6. Ni, X., Hui, C., Su, N., Jiang, W., and Liu, F. (2018). Monte Carlo simulations of electrical percolation in multicomponent thin films with nanofillers. Nanotechnology 29, 075401. 10.1088/1361-6528/aaa0be.

7. Koh, I., Rowe, H.I., and Holland, J.D. (2013). Graph and circuit theory connectivity models of conservation biological control agents. Ecol Appl 23, 1554–1573. 10.1890/12-1595.1.

8. Altmann, M. (1993). Reinterpreting network measures for models of disease transmission. Soc Networks 15, 1–17. 10.1016/0378-8733(93)90019-h.

9. McRae, B.H., Dickson, B.G., Keitt, T.H., and Shah, V.B. (2008). Using circuit theory to model connectivity in ecology, evolution, and conservation. Ecology 89, 2712–2724. 10.1890/07-1861.1.

10. Dörfler, F., and Bullo, F. (2013). Kron Reduction of Graphs With Applications to Electrical Networks. Ieee T Circuits-I 60, 150–163. 10.1109/Tcsi.2012.2215780.

11. Oughtred, R., Rust, J., Chang, C., Breitkreutz, B.J., Stark, C., Willems, A., Boucher, L., Leung, G., Kolas, N., Zhang, F., Dolma, S., Coulombe-Huntington, J., Chatr-Aryamontri, A., Dolinski, K., and Tyers, M. (2021). The BioGRID database: A comprehensive biomedical resource of curated protein, genetic, and chemical interactions. Protein Sci 30, 187–200. 10.1002/pro.3978.

12. Stark, C., Breitkreutz, B.J., Reguly, T., Boucher, L., Breitkreutz, A., and Tyers, M. (2006). BioGRID: a general repository for interaction datasets. Nucleic Acids Res 34, D535–539. 10.1093/nar/gkj109.

13. Oughtred, R., Stark, C., Breitkreutz, B.J., Rust, J., Boucher, L., Chang, C., Kolas, N., O’Donnell, L., Leung, G., McAdam, R., Zhang, F., Dolma, S., Willems, A., Coulombe-Huntington, J., Chatr-Aryamontri, A., Dolinski, K., and Tyers, M. (2019). The BioGRID interaction database: 2019 update. Nucleic Acids Res 47, D529–D541. 10.1093/nar/gky1079.

14. Gene Ontology, C. (2015). Gene Ontology Consortium: going forward. Nucleic Acids Res 43, D1049–1056. 10.1093/nar/gku1179.

15. Ashburner, M., Ball, C.A., Blake, J.A., Botstein, D., Butler, H., Cherry, J.M., Davis, A.P., Dolinski, K., Dwight, S.S., Eppig, J.T., Harris, M.A., Hill, D.P., Issel-Tarver, L., Kasarskis, A., Lewis, S., Matese, J.C., Richardson, J.E., Ringwald, M., Rubin, G.M., and Sherlock, G. (2000). Gene ontology: tool for the unification of biology. The Gene Ontology Consortium. Nat Genet 25, 25–29. 10.1038/75556.

16. Rai, S.K., Singh, D., and Sarangi, P.P. (2023). Role of RhoG as a regulator of cellular functions: integrating insights on immune cell activation, migration, and functions. Inflamm Res 72, 1453–1463. 10.1007/s00011-023-01761-9.

17. Alberti, S., and Hyman, A.A. (2021). Biomolecular condensates at the nexus of cellular stress, protein aggregation disease and ageing. Nat Rev Mol Cell Biol 22, 196–213. 10.1038/s41580-020-00326-6.

18. Banani, S.F., Lee, H.O., Hyman, A.A., and Rosen, M.K. (2017). Biomolecular condensates: organizers of cellular biochemistry. Nat Rev Mol Cell Biol 18, 285–298. 10.1038/nrm.2017.7.

19. Turowski, T.W., and Boguta, M. (2021). Specific Features of RNA Polymerases I and III: Structure and Assembly. Front Mol Biosci 8, 680090. 10.3389/fmolb.2021.680090.

20. Jacobs, R.Q., and Schneider, D.A. (2024). Transcription elongation mechanisms of RNA polymerases I, II, and III and their therapeutic implications. J Biol Chem 300, 105737. 10.1016/j.jbc.2024.105737.

21. Vannini, A., and Cramer, P. (2012). Conservation between the RNA polymerase I, II, and III transcription initiation machineries. Mol Cell 45, 439–446. 10.1016/j.molcel.2012.01.023.

22. Yeganeh, M., and Hernandez, N. (2020). RNA polymerase III transcription as a disease factor. Genes Dev 34, 865–882. 10.1101/gad.333989.119.

23. Dergai, O., and Hernandez, N. (2019). How to Recruit the Correct RNA Polymerase? Lessons from snRNA Genes. Trends Genet 35, 457–469. 10.1016/j.tig.2019.04.001.

24. Dergai, O., Cousin, P., Gouge, J., Satia, K., Praz, V., Kuhlman, T., Lhote, P., Vannini, A., and Hernandez, N. (2018). Mechanism of selective recruitment of RNA polymerases II and III to snRNA gene promoters. Genes Dev 32, 711–722. 10.1101/gad.314245.118.

25. Zentner, G.E., Saiakhova, A., Manaenkov, P., Adams, M.D., and Scacheri, P.C. (2011). Integrative genomic analysis of human ribosomal DNA. Nucleic Acids Res 39, 4949–4960. 10.1093/nar/gkq1326.

26. Cheng, R., Rajendra, K.C., Lizarazo, S., Yuan, J., Zhou, S., and Van Bortle, K. (2025). The NuRD complex shapes RNA polymerase III activity at highly expressed tRNA gene clusters, tuning the dynamic range of cellular tRNA pools. bioRxiv, 2025.2012.2012.694012. 10.64898/2025.12.12.694012.

27. Engel, C., Plitzko, J., and Cramer, P. (2016). RNA polymerase I-Rrn3 complex at 4.8 A resolution. Nat Commun 7, 12129. 10.1038/ncomms12129.

28. Jochem, L., Ramsay, E.P., and Vannini, A. (2017). RNA polymerase I, bending the rules? EMBO J 36, 2664–2666. 10.15252/embj.201797924.

29. Richter, W.F., Nayak, S., Iwasa, J., and Taatjes, D.J. (2022). The Mediator complex as a master regulator of transcription by RNA polymerase II. Nat Rev Mol Cell Biol 23, 732–749. 10.1038/s41580-022-00498-3.

30. Hsieh, Y.J., Wang, Z., Kovelman, R., and Roeder, R.G. (1999). Cloning and characterization of two evolutionarily conserved subunits (TFIIIC102 and TFIIIC63) of human TFIIIC and their involvement in functional interactions with TFIIIB and RNA polymerase III. Mol Cell Biol 19, 4944–4952. 10.1128/MCB.19.7.4944.

31. Birch, J.L., Tan, B.C., Panov, K.I., Panova, T.B., Andersen, J.S., Owen-Hughes, T.A., Russell, J., Lee, S.C., and Zomerdijk, J.C. (2009). FACT facilitates chromatin transcription by RNA polymerases I and III. EMBO J 28, 854–865. 10.1038/emboj.2009.33.

32. Carre, C., and Shiekhattar, R. (2011). Human GTPases associate with RNA polymerase II to mediate its nuclear import. Mol Cell Biol 31, 3953–3962. 10.1128/MCB.05442-11.

33. Graczyk, D., Debski, J., Muszynska, G., Bretner, M., Lefebvre, O., and Boguta, M. (2011). Casein kinase II-mediated phosphorylation of general repressor Maf1 triggers RNA polymerase III activation. Proc Natl Acad Sci U S A 108, 4926–4931. 10.1073/pnas.1010010108.

34. Hu, P., Samudre, K., Wu, S., Sun, Y., and Hernandez, N. (2004). CK2 phosphorylation of Bdp1 executes cell cycle-specific RNA polymerase III transcription repression. Mol Cell 16, 81–92. 10.1016/j.molcel.2004.09.008.

35. Hu, P., Wu, S., and Hernandez, N. (2003). A minimal RNA polymerase III transcription system from human cells reveals positive and negative regulatory roles for CK2. Mol Cell 12, 699–709. 10.1016/j.molcel.2003.08.011.

36. Johnston, I.M., Allison, S.J., Morton, J.P., Schramm, L., Scott, P.H., and White, R.J. (2002). CK2 forms a stable complex with TFIIIB and activates RNA polymerase III transcription in human cells. Mol Cell Biol 22, 3757–3768. 10.1128/MCB.22.11.3757-3768.2002.

37. Steinhoff, J.S., Lass, A., and Schupp, M. (2021). Biological Functions of RBP4 and Its Relevance for Human Diseases. Front Physiol 12, 659977. 10.3389/fphys.2021.659977.

38. Muzi-Falconi, M., Giannattasio, M., Foiani, M., and Plevani, P. (2003). The DNA polymerase alpha-primase complex: multiple functions and interactions. ScientificWorldJournal 3, 21–33. 10.1100/tsw.2003.05.

39. Pellegrini, L. (2012). The Pol alpha-primase complex. Subcell Biochem 62, 157–169. 10.1007/978-94-007-4572-8_9.

40. Ruepp, A., Brauner, B., Dunger-Kaltenbach, I., Frishman, G., Montrone, C., Stransky, M., Waegele, B., Schmidt, T., Doudieu, O.N., Stumpflen, V., and Mewes, H.W. (2008). CORUM: the comprehensive resource of mammalian protein complexes. Nucleic Acids Res 36, D646–650. 10.1093/nar/gkm936.

41. Tsitsiridis, G., Steinkamp, R., Giurgiu, M., Brauner, B., Fobo, G., Frishman, G., Montrone, C., and Ruepp, A. (2023). CORUM: the comprehensive resource of mammalian protein complexes-2022. Nucleic Acids Res 51, D539–D545. 10.1093/nar/gkac1015.

42. Lai, A.Y., and Wade, P.A. (2011). Cancer biology and NuRD: a multifaceted chromatin remodelling complex. Nat Rev Cancer 11, 588–596. 10.1038/nrc3091.

43. Xue, Y., Wong, J., Moreno, G.T., Young, M.K., Cote, J., and Wang, W. (1998). NURD, a novel complex with both ATP-dependent chromatin-remodeling and histone deacetylase activities. Mol Cell 2, 851–861. 10.1016/s1097-2765(00)80299-3.

44. Gagliardi, M., Strazzullo, M., and Matarazzo, M.R. (2018). DNMT3B Functions: Novel Insights From Human Disease. Front Cell Dev Biol 6, 140. 10.3389/fcell.2018.00140.

45. Piunti, A., and Shilatifard, A. (2021). The roles of Polycomb repressive complexes in mammalian development and cancer. Nat Rev Mol Cell Biol 22, 326–345. 10.1038/s41580-021-00341-1.

46. Lizarazo, S., Chen, E., K C, R., and Van Bortle, K. (2026). dominatR: Feature Dominance-based R Package for Genomic Data (Bioconductor).

47. Klein, D.J., and Randic, M. (1993). Resistance Distance. J Math Chem 12, 81–95. DOI 10.1007/Bf01164627.

48. Wu, T., Hu, E., Xu, S., Chen, M., Guo, P., Dai, Z., Feng, T., Zhou, L., Tang, W., Zhan, L., Fu, X., Liu, S., Bo, X., and Yu, G. (2021). clusterProfiler 4.0: A universal enrichment tool for interpreting omics data. Innovation (Camb) 2, 100141. 10.1016/j.xinn.2021.100141.

49. Ju, W.Y., Li, J.X., Yu, W.R., and Zhang, R.C. (2016). iGraph: an incremental data processing system for dynamic graph. Front Comput Sci-Chi 10, 462–476. 10.1007/s11704-016-5485-7.

50. Gu, Y., Wang, C.L., Ma, J., Nemiroff, R.J., and Kao, D.L. (2015). iGraph: A Graph-Based Technique for Visual Analytics of Image and Text Collections. Proc Spie 9397. Artn 939708 10.1117/12.2074198.

51. Csardi, G., and Nepusz, T. (2006). The igraph software package for complex network research. InterJournal, Complex Systems 1695, 1–9.

52. Ahmad, S., Jose da Costa Gonzales, L., Bowler-Barnett, E.H., Rice, D.L., Kim, M., Wijerathne, S., Luciani, A., Kandasaamy, S., Luo, J., Watkins, X., Turner, E., Martin, M.J., and UniProt, C. (2025). The UniProt website API: facilitating programmatic access to protein knowledge. Nucleic Acids Res 53, W547–W553. 10.1093/nar/gkaf394.

53. UniProt, C. (2025). UniProt: the Universal Protein Knowledgebase in 2025. Nucleic Acids Res 53, D609–D617. 10.1093/nar/gkae1010.

54. Smedley, D., Haider, S., Durinck, S., Pandini, L., Provero, P., Allen, J., Arnaiz, O., Awedh, M.H., Baldock, R., Barbiera, G., Bardou, P., Beck, T., Blake, A., Bonierbale, M., Brookes, A.J., Bucci, G., Buetti, I., Burge, S., Cabau, C., Carlson, J.W., Chelala, C., Chrysostomou, C., Cittaro, D., Collin, O., Cordova, R., Cutts, R.J., Dassi, E., Di Genova, A., Djari, A., Esposito, A., Estrella, H., Eyras, E., Fernandez-Banet, J., Forbes, S., Free, R.C., Fujisawa, T., Gadaleta, E., Garcia-Manteiga, J.M., Goodstein, D., Gray, K., Guerra-Assuncao, J.A., Haggarty, B., Han, D.J., Han, B.W., Harris, T., Harshbarger, J., Hastings, R.K., Hayes, R.D., Hoede, C., Hu, S., Hu, Z.L., Hutchins, L., Kan, Z., Kawaji, H., Keliet, A., Kerhornou, A., Kim, S., Kinsella, R., Klopp, C., Kong, L., Lawson, D., Lazarevic, D., Lee, J.H., Letellier, T., Li, C.Y., Lio, P., Liu, C.J., Luo, J., Maass, A., Mariette, J., Maurel, T., Merella, S., Mohamed, A.M., Moreews, F., Nabihoudine, I., Ndegwa, N., Noirot, C., Perez-Llamas, C., Primig, M., Quattrone, A., Quesneville, H., Rambaldi, D., Reecy, J., Riba, M., Rosanoff, S., Saddiq, A.A., Salas, E., Sallou, O., Shepherd, R., Simon, R., Sperling, L., Spooner, W., Staines, D.M., Steinbach, D., Stone, K., Stupka, E., Teague, J.W., Dayem Ullah, A.Z., Wang, J., Ware, D., Wong-Erasmus, M., Youens-Clark, K., Zadissa, A., Zhang, S.J., and Kasprzyk, A. (2015). The BioMart community portal: an innovative alternative to large, centralized data repositories. Nucleic Acids Res 43, W589–598. 10.1093/nar/gkv350.

55. Durinck, S., Moreau, Y., Kasprzyk, A., Davis, S., De Moor, B., Brazma, A., and Huber, W. (2005). BioMart and Bioconductor: a powerful link between biological databases and microarray data analysis. Bioinformatics 21, 3439–3440. 10.1093/bioinformatics/bti525.

56. Verfaillie, A., Imrichova, H., Janky, R., and Aerts, S. (2015). iRegulon and i-cisTarget: Reconstructing Regulatory Networks Using Motif and Track Enrichment. Curr Protoc Bioinformatics 52, 2 16 11-12 16 39. 10.1002/0471250953.bi0216s52.

57. Imrichova, H., Hulselmans, G., Atak, Z.K., Potier, D., and Aerts, S. (2015). i-cisTarget 2015 update: generalized cis-regulatory enrichment analysis in human, mouse and fly. Nucleic Acids Res 43, W57–64. 10.1093/nar/gkv395.

58. Herrmann, C., Van de Sande, B., Potier, D., and Aerts, S. (2012). i-cisTarget: an integrative genomics method for the prediction of regulatory features and cis-regulatory modules. Nucleic Acids Res 40, e114. 10.1093/nar/gks543.

59. Cantarella, S., Neffe, J., Hermes, M., Rauscher, F., Schwarz, R., Jaisankar, P., Schafer, E., and Caudron-Herger, M. (2026). RBP2GO 2.0: Integrating disease associations and sequence features to explore RNA-binding protein functions. Nucleic Acids Res 54, D481–D488. 10.1093/nar/gkaf1258.

60. Caudron-Herger, M., Jansen, R.E., Wassmer, E., and Diederichs, S. (2021). RBP2GO: a comprehensive pan-species database on RNA-binding proteins, their interactions and functions. Nucleic Acids Res 49, D425–D436. 10.1093/nar/gkaa1040.

