## Supplementary figures and images for "Reciprocal effective conductance-based inference of protein-protein relationships delineates protein network hubs and macromolecular properties"

### Supplementary Video 1

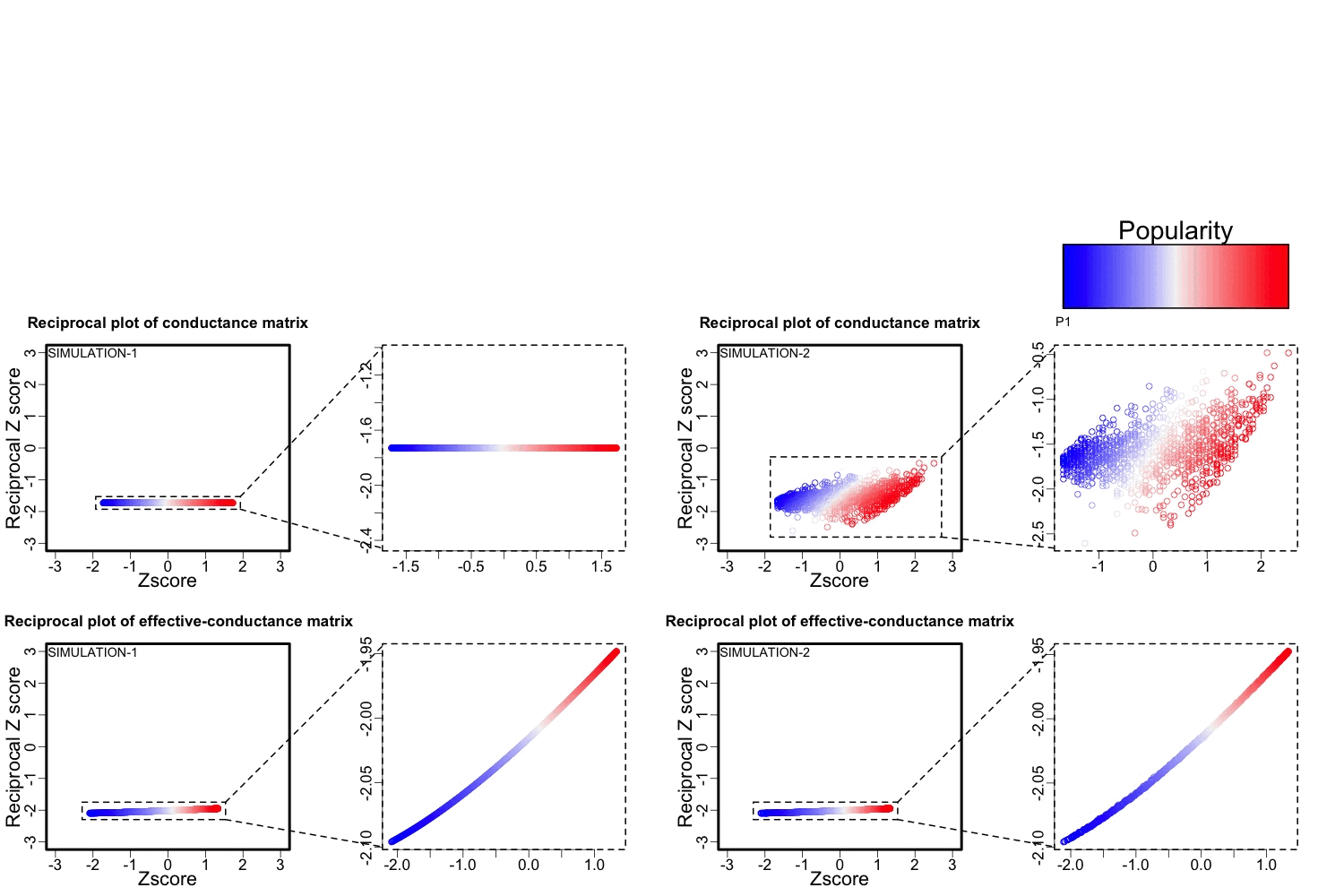

### Supplementary Video 2

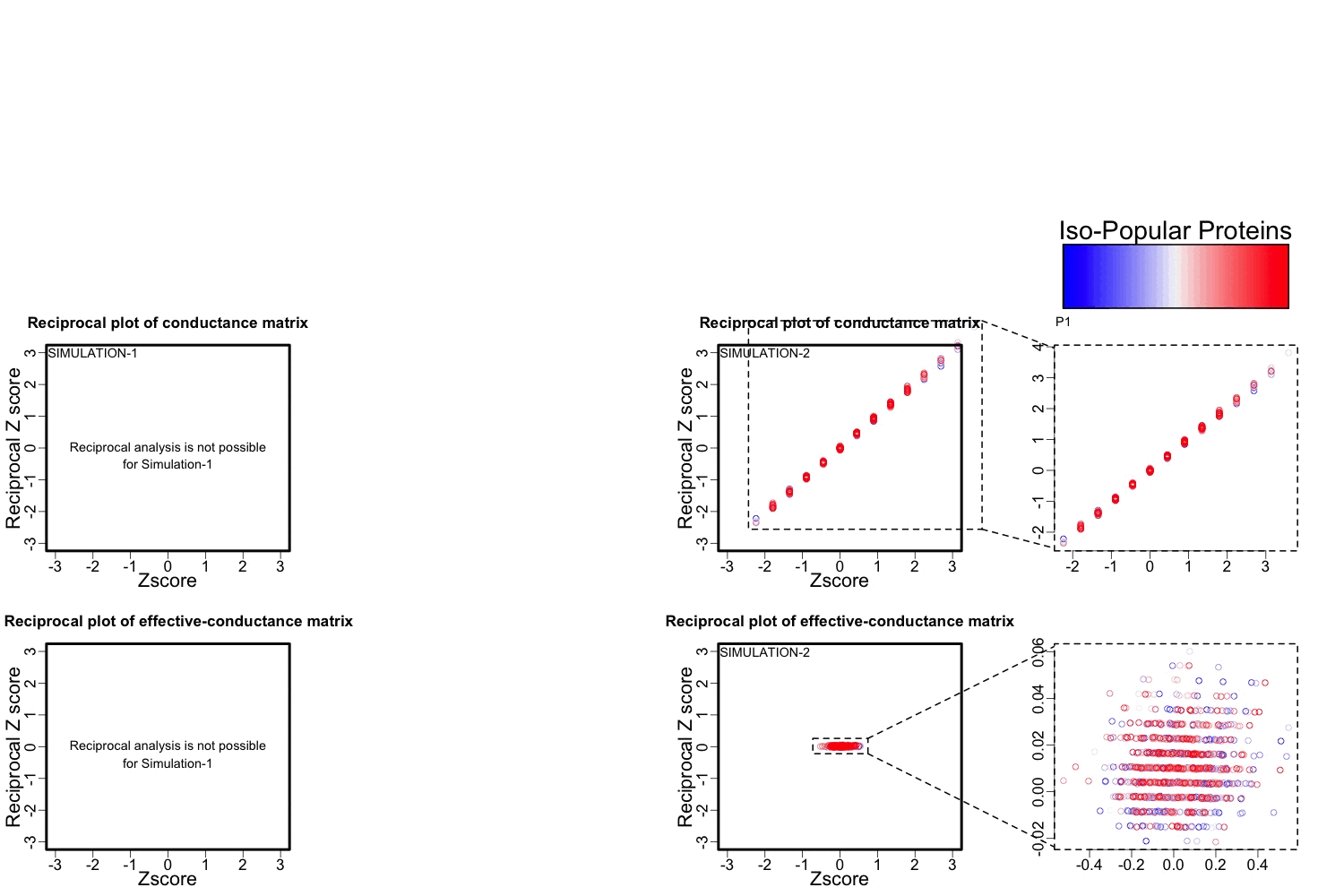
